# Brain solute transport is more rapid in periarterial than perivenous spaces

**DOI:** 10.1101/2021.03.23.436588

**Authors:** Vegard Vinje, Erik NTP Bakker, Marie E Rognes

## Abstract

**Background:** Perivascular fluid flow, of cerebrospinal or interstitial fluid in spaces surrounding brain blood vessels, is recognized as a key component underlying brain transport and clearance. An important open question is how and to what extent differences in vessel type or geometry affect perivascular fluid flow and transport.

**Methods:** Using computational modelling in both idealized and image-based geometries, we study and compare fluid flow and solute transport in pial (surface) periarterial and perivenous spaces.

**Results:** Our findings demonstrate that differences in geometry between arterial and venous pial perivascular spaces (PVSs) lead to higher net CSF flow, more rapid tracer transport and earlier arrival times of injected tracers in periarterial spaces compared to perivenous spaces.

**Conclusions:** These findings can explain the experimentally observed rapid appearance of tracers around arteries, and the delayed appearance around veins without the need of a circulation through the parenchyma, but rather by direct transport along the PVSs.

---

Perivascular fluid flow, of cerebrospinal or interstitial fluid in spaces surrounding brain blood vessels, is recognized as a key component underlying brain transport and clearance [1, 2, 3, 4, 5]. In spite of its potential importance, several aspects of perivascular fluid flow are poorly characterized. In particular, the shape and structure of perivascular spaces (PVSs), as well as differences between periarterial and perivenous spaces and flow, remain enigmatic and disputed.

Traditionally, PVSs at the brain surface (pial PVSs) have been described and represented as annular structures surrounding the vasculature, either disjoint from the sub-arachnoid space (SAS) [6, 7, 3, 8], or as widenings of the SAS with reduced flow resistance [9]. When observing injected tracers in the murine brain via in-vivo two-photon imaging and fluorescence microscopy, the tracer is predominantly found along pial arteries and arterioles [7, 10]. Careful examination of such images [11, Figure 1D] indicates that after an initial spread of tracer along arteries, the tracer is later present over the entire surface of the brain, thus suggesting tracer movement from the pial PVS into the adjacent SAS. The PVSs around arteries also appear to connect to the PVSs of veins, either via the SAS or directly where arteries and veins cross [11, Figure1]. These observations correspond well with other postmortem confocal images in which the tracer signal in the PVSs of arteries, veins, and SAS were continuous [9].

**Figure 1.**
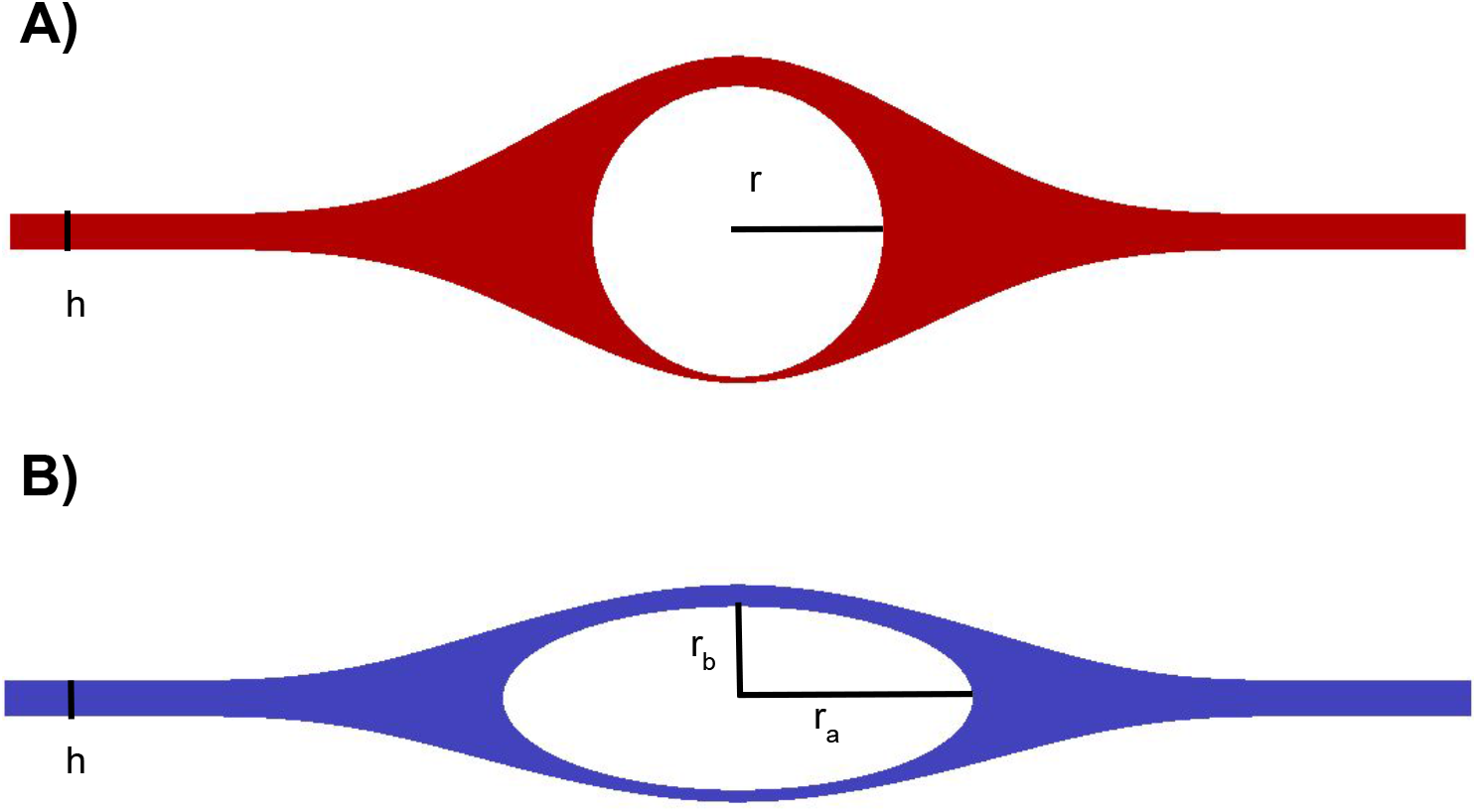
Computational domains for the idealized pial perivascular spaces surrounding an artery (A) and a vein (B) and simulating Stokes flow. In the baseline models, *r* = 20*µ*m, *ra* = 32*µ*m, *rb* = 12.5*µ*m and *h* = 5*µ*m.

**Figure 2.**
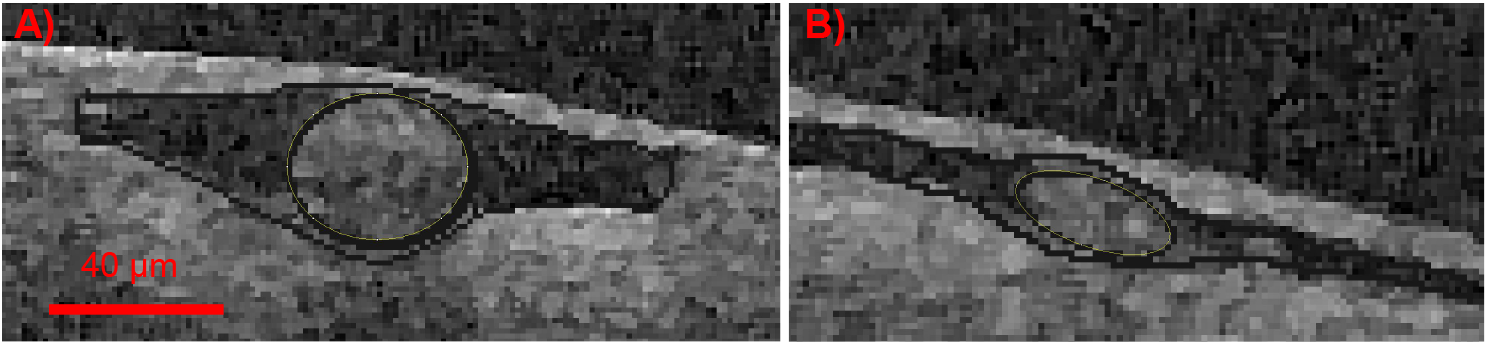
Blood vessels from OCT surrounded by PVS/SAS were used to create image-based geometries for the artery (A) and the vein (B). Scale bar indicates dimensions after scaling.

Perivascular fluid flow around arteries has now been studied extensively, while less evidence exists regarding the role and characteristics of perivenous flow. In and around large arteries on the brain surface, it seems well documented that the bulk fluid flow direction follows the blood flow direction [7, 10]. Perivenous spaces have been suggested as exit routes, draining interstitial fluid and solutes out of the brain and the brain environment [12]. Evidence for this view includes later arrival of intracisternally injected tracer to venous PVSs compared to arterial PVSs, and tracer accumulation around venous PVSs after intraparenchymal injection [3]. However, in earlier work, it was suggested that the direction of PVS fluid flow varies in a more unpredictable manner [2].

The anatomy, shape, and structure of perivascular spaces may vary by location and change over time [5]. In particular, there may be persistent and systematic differences between the spaces surrounding pial arteries and veins [9]. A key open question is how and to what extent differences in shape affect PVS fluid flow and transport. Several theoretical and computational models have used the space between two concentric cylinders to model the PVS [13, 14, 15, 16, 17]. Mestre et al. [7] have suggested, based on live tracer studies, that PVSs may be non-connecting, with two disjoint compartments forming on each side of pial arteries. Folowing these observations, Tithof et al. [18] considered a confined PVS separated from the SAS, and found that an elliptic PVS was optimal for fluid transport given a constraint on the cross sectional area. An elliptical model with two disjoint PVSs was also assumed by Kedarasetti et al. to study perivascular pumping [19]. However, to the best of our knowledge, the physiological implications of differences in shape on periarterial and perivenous flow and solute transport have not been adequately characterized. Perivenous flow within the brain parenchyma has been studied in a 1D-model by Faghih and Sharp [20], but models of the perivenous space at a level of detail comparable to the modeling studies mentioned above [13, 14, 15, 16, 19] is lacking.

In this paper, using computational modelling in both idealized and image-based geometries, we study and compare fluid flow and solute transport in pial (surface) periarterial and perivenous spaces. We first systematically address how PVS fluid velocities change with differences in arterial and venous PVS geometries. Next, we predict arrival differences of tracer in perivascular spaces based on differences in fluid velocities. Our findings demonstrate that fluid flow and solute transport is more rapid in spaces of periarterial shape when compared to spaces of perivenous shape, and that these geometrical differences alone predict a delayed tracer arrival in pial perivenous spaces.

## Methods

### Domains and meshes

We studied families of three-dimensional geometries Ω of pial PVSs: (a) families of idealized PVSs around an artery (A0) and/or a vein (V0), and (b) an image-based set of PVSs surrounding an artery (A1) and/or a vein (V1) extracted from human optical coherence tomography (OCT) images. To compare with experimental data of tracer transport, all vessels and PVSs were scaled to the mouse size (such that *r* = 20*µ*m for the arteries in agreeement with [7]). Cross-sections of the periarterial and perivenous spaces are illustrated in Figures 1 (idealized) and 2 (image-based); these cross-sections were extruded by 5 mm along the third axis to define the computational domains.

Each cross-section of the idealized periarterial spaces was assumed to surround a circular artery and to be continuous with the SAS with more open space on each side of the vessel than above and below [9]. The outer PVS surfaces were defined by spline curves via sets of control points (for .geo-files, see the data set [21]). The minimal SAS height (h)was set to 5 *µ*m, the maximal SAS height (including the artery with diameter 2r = 40 *µ*m) was set to 45 *µ*m, and the PVS width representing the distance between the arterial surface and the end of the PVS tapering was set to 40 *µ*m. For the idealized perivenous spaces, the vein was assumed to form an ellipse with major axis *r_a_* = 32 *µ*m and minor axis *r_b_* = 12.5 *µ*m. The cross-sectional area of the artery and the vein are thus equal. In the narrow SAS region, the perivenous height was comparable to that of the artery (h), while the maximal SAS height (including the vein) was 30 *µ*m.

To systematically quantify the effect of PVS geometry on maximal PVS flow velocity, we created two further families of computational domains parametrized by (i) a vascularity *ρ* ∈ [0, 1] (with *ρ* = 0 and *ρ* = 1 corresponding to the venous and arterial PVS, respectively, and *ρ* ∈ (0, 1) linearly transforming between these two geometries) and (ii) the periarterial width *w* ∈ (30, 70)*µ*m. For both families, we uniformly sampled 100 meshes. The gradual changes in geometry for the two cases are visualized in Figure 5 and Additional Files 1–2.

The image-based geometries were constructed from previously published OCT data [10] in which the PVSs surrounding two vessels were easily recognizable and identifiable, and meshed via Gmsh [22].

### Computational fluid dynamics in the pial PVS

In the pulsatile perivascular flow observed experimentally [7, 10], flow amplitudes are similar compared to the net flow (≈10 *µ*m/s vs ≈20 *µ*m/s). The dominant transport mechanism over time will thus be the net movement of fluid, with only minor effects of mixing [23]. Therefore, together with the assumptions that pial PVSs are continuous with the SAS [9] and open [24], we modelled the convective flow in the PVS as an incompressible Stokes flow; that is, the fluid velocity *v* = *v* (*x, y, z*) and pressure *p* = *p*(*x, y, z*) for (*x, y, z*) ∈ Ω solve

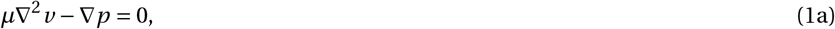

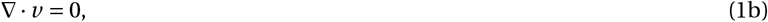

under additional given boundary conditions.

In the extruded domains considered here, in which each cross-section of each geometry is constant, the incompressible Stokes equations subject to a constant pressure difference over the domain length can be reduced to the following equation for the axial velocity *v* = *v* (*x*, *y*) (velocity in the z-direction) over the two-dimensional cross-sections:

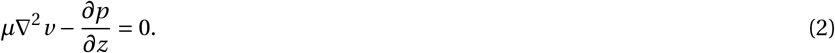

We let the flow be driven by a constant pressure gradient 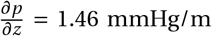. Even though such a gradient has only been reported as a pulsatile gradient [25], static gradients of this magnitude have been shown to create net velocities of around 20 *µ*m/s in the PVS [16] in agreement with experimental observations of injected microspheres in pial PVS. We also conducted experiments with a lower pressure gradient: 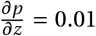 mmHg/m [25] corresponding to the third circulation’s production of CSF alone [26]. On the vessel wall, and on the upper (dura mater) and lower (pia mater) walls we assumed a no slip condition (i.e. *v* = 0), while on the side walls (extending further in the SAS), we used the natural Neumann (symmetry) condition (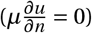).

### Convection-diffusion modelling

We simulated the distribution and evolution of a solute (e.g. tracer or microspheres) in the pial PVS in response to a steady injection at one end (*z* = 0) of the PVS geometries. We assume that the solute concentration *u* = *u*(*x*, *y*, *z*, *t*) ∈ Ω will be governed by the convection-diffusion system:

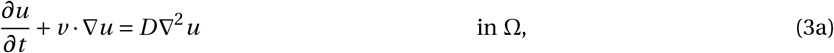

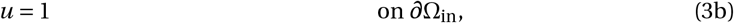

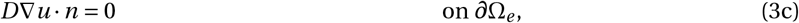

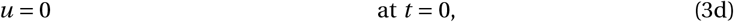

where *v* is the convective fluid velocity, and *D* is the diffusion coefficient in water for the given solute. We denote the inlet end of the geometry by *∂*Ω_in_, while *∂*Ω_*e*_ is the remaining part of the domain boundary *∂*Ω. The boundary and initial conditions (3b) - (3d) describe a constant injection at one end of the PVS geometry with no initial tracer anywhere in the PVS or SAS. To model pure diffusion in the PVS, we set *v* = 0. When including convection, we first solved the pressure-driven Stokes Equation (2) to find the fluid velocity *v* = *v* (*x*, *y*) = *v* (*x*, *y*, *z*).

### Material parameters

We set the CSF viscosity *µ* = 0.7 Pa s [16]. The diffusion coefficient *D* varies by molecular size, and small particles typically diffuse faster than larger particles. We set the diffusion coefficient to *D* = 1.0 × 10^−8^ cm^2^/s to represent unhindered diffusion in the CSF-filled PVS of the large (2000 kDa) tracer used for volumetric imaging by Mestre et al [7]. This estimate is based on reported apparent diffusion coefficients *D*\* = *Dλ*^2^ for 70 kDa Dextran of 0.84 × 10^−7^ cm^2^/s in brain neuropil with a tortuosity *λ* of 1.6 [27] (thus *D* = 2.15 × 10^−7^ cm^2^/s), and 5.3×10^−7^ cm^2^/s (thus *D* = 1.36×10^−6^ cm^2^/s) for the smaller 3 kDa Texas Red dextran [3, 28]; and the reported apparent diffusion coefficients in articular cartilege of approximately 7.5 × 10^−7^ cm^2^/s for 3 kDa, 3.0 × 10^−7^ cm^2^/s for 70 kDa, and 4.0 × 10^−8^ cm^2^/s for 500 kDa dextran [29]. Microspheres of approximately 1 *µ*m in diameter [7] are larger than these substances, and would be expected to diffuse even slower. In addition to the small diffusion coefficient for the 2000 kDa tracer, we performed simulations using the ≈100x higher diffusion coefficient of 3 kDa Texas Red dextran as well.

### Quantities of interest

The concentration *u* is a normalized value (between 0 and 1) and set to 1 at the injection site (*z* = 0). As our primary quantity of interest, we measure the average tracer concentration 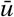, averaged over the cross-section *∂*Ω_end_ at the end of the domain (where *z* = *L*), over time:

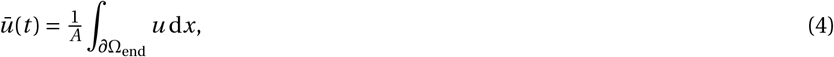

where *A* is the area of *∂*Ω_end_. In addition to 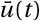, we report the velocity and concentration fields in the periarterial and perivenous spaces.

### Numerical solution and verification

The partial differential equations were solved using the finite element method and the FEniCS software suite [30]. The reduced Stokes equations (2) were solved using a first order (continuous piecewise linear) approximation (in space). The convection-diffusion equations (3) were solved using a first order approximation in space and time (implicit Euler scheme) with a uniform time step ∆*t* = 10 s and mesh resolution ∆*x* = 1 *µ*m until *T* = 120 minutes. The discrete velocity field *v* = *v* (*x*, *y*) was interpolated onto each vertex of the three-dimensional mesh: *v* (*x*, *y*, *z*) = *v* (*x*, *y*). For the pure diffusion simulations, we used a time step of ∆*t* = 1 day and an end time of *T* = 150 days. Numerical convergence tests were performed to ensure that the reported results were independent of mesh resolution and the choice of time step (see Additional File 3). The resolution of the mesh used for simulations are shown in Additional File 4.

## Results

In all experiments tracer spread from the site of injection towards the site of measurement (See Figure 3A, B). The rate of transport of tracer was dominated by convection and varied between different geometries.

**Figure 3.**
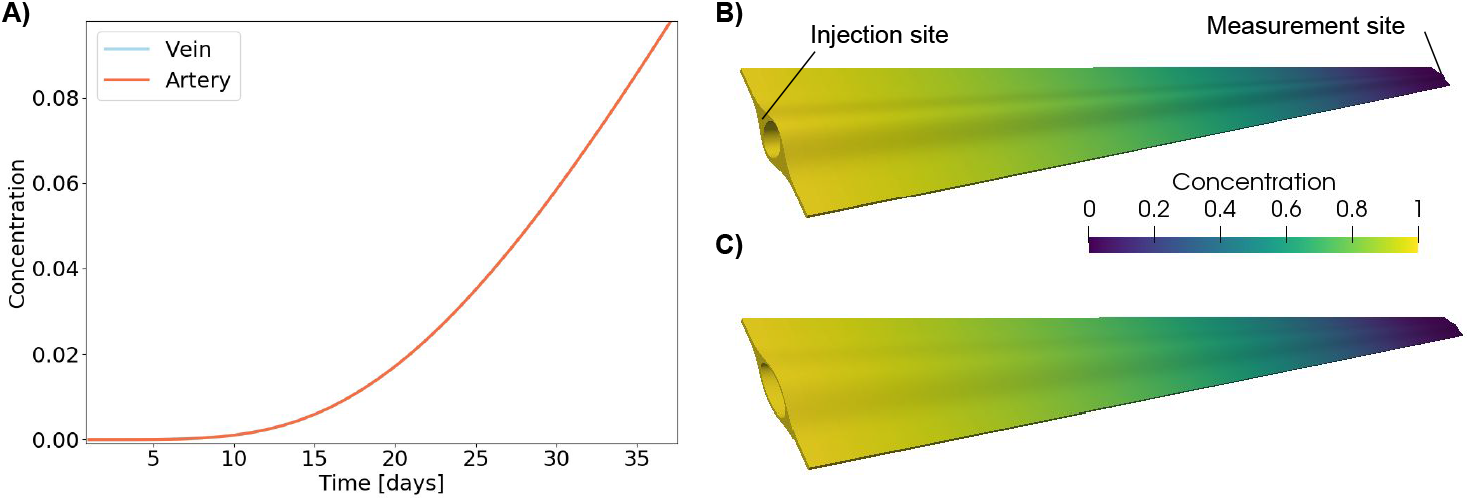
Diffusion of large tracers in PVS occurs on the time scale of days. In A) the concentration of tracer at the point of measurement is shown for the artery and the vein. The two graphs are identical, showing that diffusion occurs at a similar rate in the two geometries. Tracer distribution after 20 days of diffusive transport is shown for the artery (B) and the vein (C).

### Diffusive transport is too slow to explain tracer movement along PVS

Even when measured only 5 mm from the injection site, diffusion was too slow to transport the large tracers of interest within the time frame (a few minutes) observed in experiments [7]: it took up to 10 days for an increase in tracer concentration to be detectable at the PVS outlets (Figure 3A). After 20 days, the average concentration at the outlet was 1-2 % of the concentration at the injection site. Moreover, diffusion transported the tracer along the artery and along the vein at the same pace (Figure 3). Figure 3B and C show the distribution along the artery (B) and the vein (C) after 20 days of diffusive transport. The tracer was evenly distributed at all cross-sections. Thus, for pure diffusion, the shape of the PVS did not influence the tracer transport.

### Periarterial geometries induce higher fluid velocities

The imposed pressure gradients induced fluid velocities in all geometries, with greater velocities in the larger open spaces surrounding the arteries (both idealized and image-based) than in other regions (see Figure 4). The peak (and average) fluid velocity in the idealized periarterial and perivenous spaces was 11 (3) and 5 (1) *µ*m/s, respectively, and 17 (6) and 3 (1) *µ*m/s in the image-based periarterial and perivenous spaces. The velocities were higher on each side of the blood vessels, with little or no flow over the top or below the vessel. We note that even though the PVS forms an open space around the artery continuous with the SAS, the PVS flow pattern generated two distinct regions with higher fluid velocities. In the SAS far away from the vessel, the maximal fluid velocity was steady at ≈1 *µ*m/s in the idealized artery and vein models. Using the maximal fluid velocities *u* reported above, and the characteristic length *L* = 5 mm, the Peclet numbers 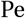 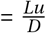 ranged from Pe = 85000 (A1) to Pe = 110 (V1 with a 100-fold increase in diffusion coefficient).

**Figure 4.**
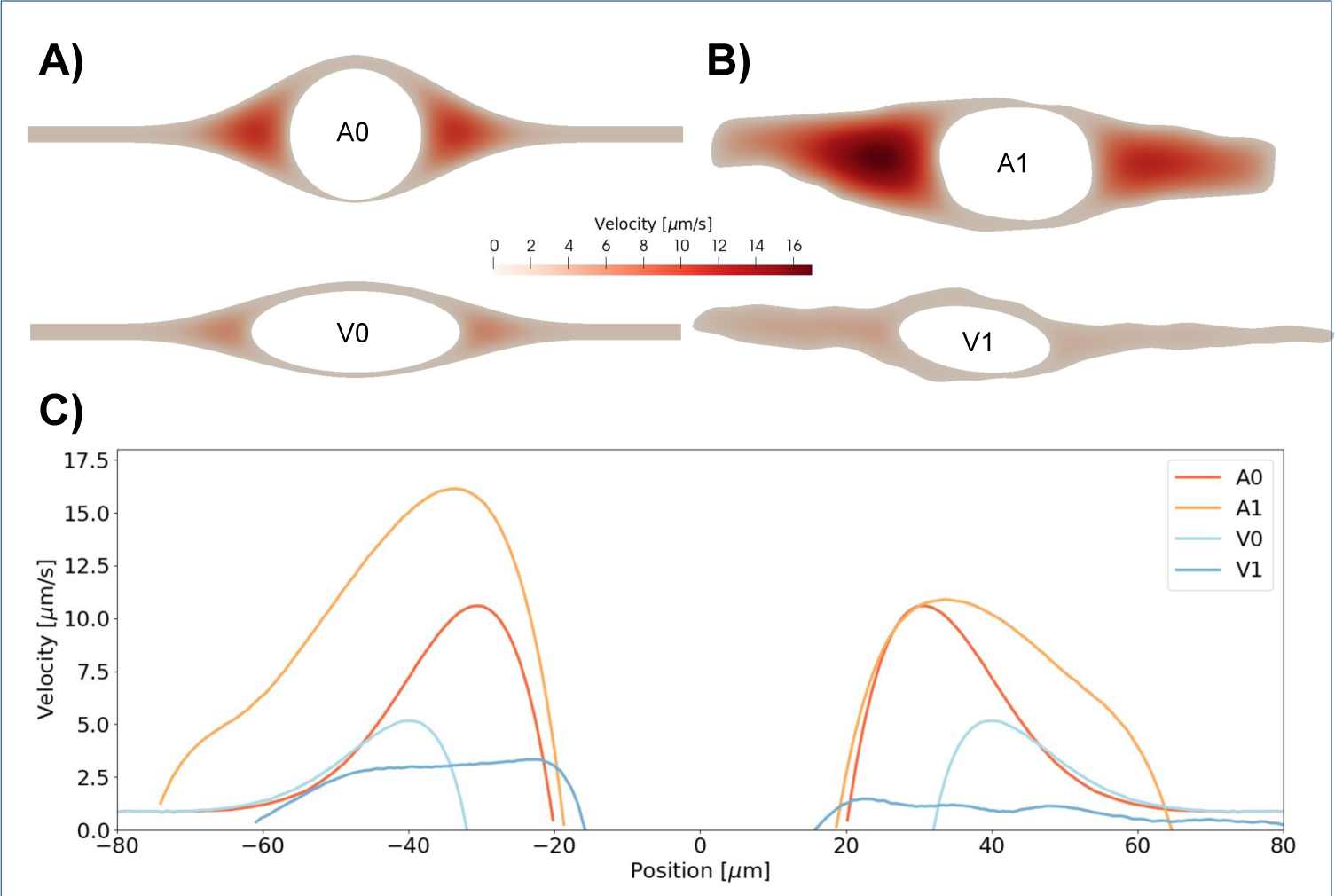
Velocity in the PVS around an artery and a vein in the idealized (A) and extracted (B) geometries. In these models, velocity has only one non-zero component, along the PVS. In both sets of geometries, flow is larger around arteries. In the idealized artery, PVS velocities reached 11 *µ*m/s in the largest open regions. In the idealized vein, fluid velocities reach 5 *µ*m/s. Flow between the pial surface and the vein as well as flow between the dura mater and the vein were negligible due to the small widths between these structures. In the image-based geometries PVS velocities reached 17 *µ*m/s around the artery and 3 *µ*m/s around the vein. C) PVS velocity as a function of position along the horizontal centerline. *x* = 0 corresponds to the center of the vessel.

With the pressure gradient corresponding to CSF production alone (according to the third circulation), the peak fluid velocity was 0.1 *µ*m/s in the idealized artery, approximately 100 times lower than in the other experiments and would not be able to explain rapid movement of tracer in the PVS (data not shown). In the geometry with the lowest convective velocity (the image-based perivenous space, V1), transport of 3 kDa Texas Red dextran was still dominated by convection. The 100-fold increase in diffusion coefficient accelerated transport by 11-12 minutes, and in particular the tracer concentration at the measurement site exceeded a threshold of 0.7 after 75 vs 87 minutes.

### Perivascular flow velocities vary linearly with PVS vascularity and width

To determine the main geometrical parameters affecting the perivascular flow, we examined the peak perivascular fluid velocity resulting from a gradual change from the idealized periarterial geometry to the idealized perivenous geometry, and from changing PVS widths (see Methods). Linear transitions between these parameterized geometries resulted in near linear changes in peak perivascular flow velocities (Figure 5,Supplementary Videos S1-S2). An increase in PVS width from 40 *µ*m (default periarterial geometry) to 70 *µ*m increased PVS velocities from 11 (3) *µ*m/s to 23 (9) *µ*m/s, while a reduction to 30 *µ*m PVS width resulted in PVS velocities of 7 (2) *µ*m/s. Thus, a narrow periarterial space generated similar perivascular flow velocities as the idealized perivenous space (7 (2) *µ*m/s vs 5 (1) *µ*m/s). We note that the linear increase observed is in contrast to the quadratic relationship between radius and maximal flow velocity in a cylinder.

**Figure 5.**
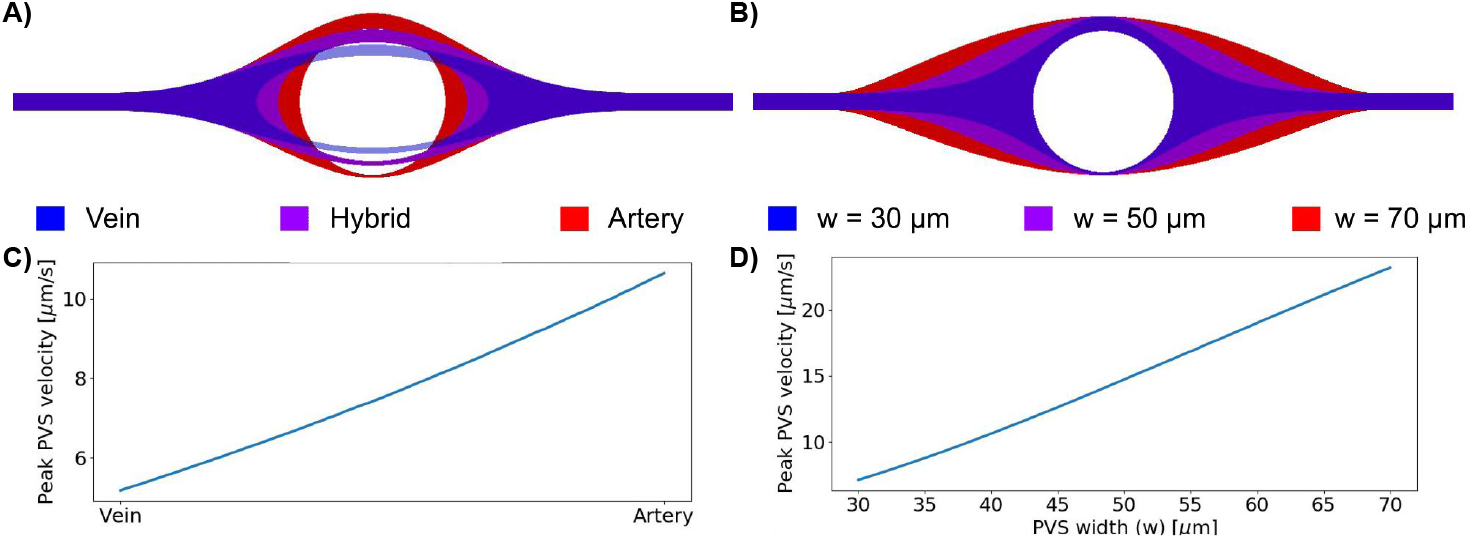
The effect of geometry change on maximal PVS velocity. A) shows a gradual change from a venous to an arterial PVS. B) shows change in PVS width for the arterial geometry. Maximal PVS velocities as a function of linear change in geometry is shown for the change from artery to vein (C), and changes in PVS width (D).In both cases the maximal PVS velocity increased linearly with changes in the geometry.

### Tracer appears first around arteries, then around veins

Simulations of convective and diffusive transport via (3) in the perivascular spaces revealed substantial differences between the periarterial and perivenous transport characteristics and the tracer arrival times at the measurement site (Figure 6).

**Figure 6.**
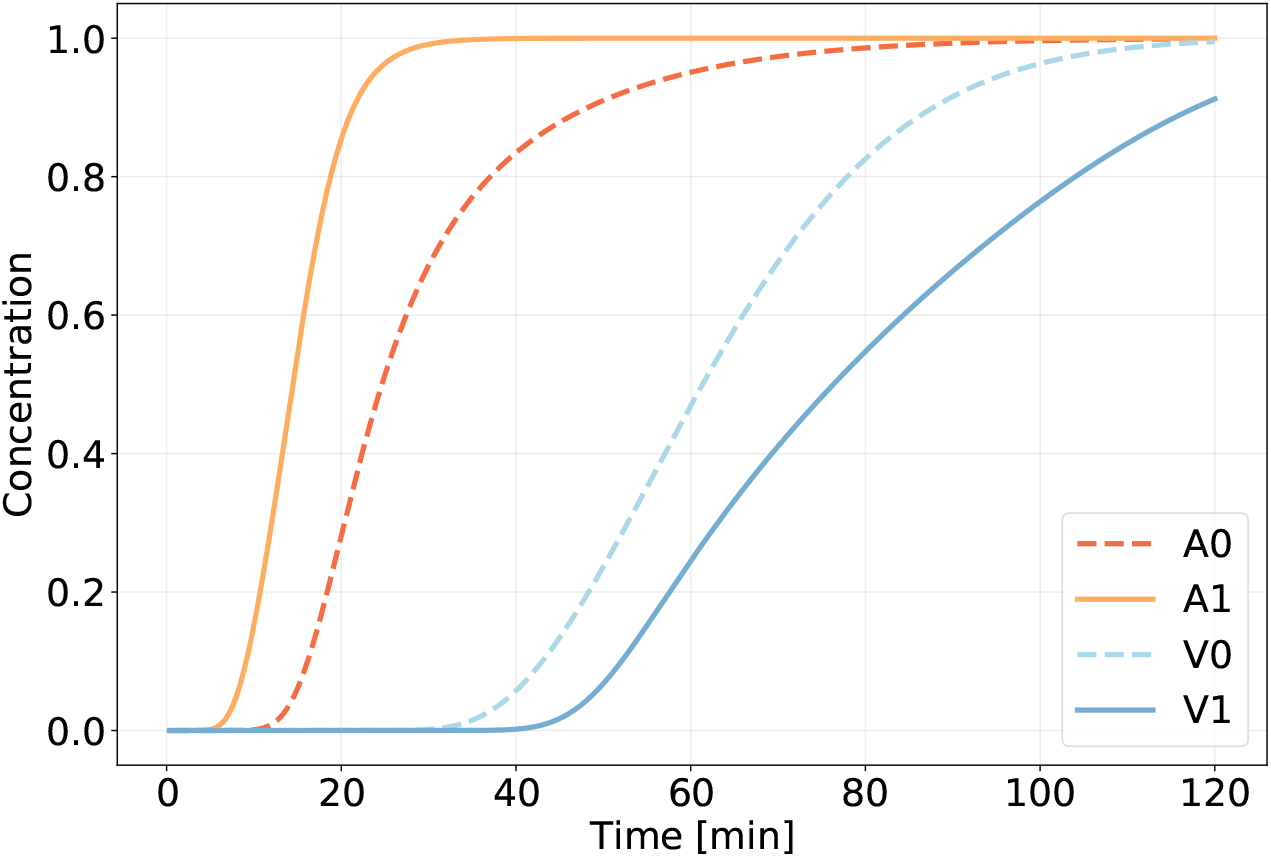
Time plots of average concentrations at the point of measurement in the idealized artery (A0) and vein PVS (V0) together with the average concentrations in the image-based artery (A1) and vein (V1) PVS. The difference between arterial and venous PVS transport is larger in the image-based geometries, but there is a significant delay in venous PVS compared to arterial PVS in both cases.

In the periarterial spaces, the average tracer concentration at the measurement site rapidly grew after 5–20 minutes, with the most rapid transport and time to peak concentration (≈25 min) in the image-based periarterial spaces. In the perivenous spaces, the measured concentration remained low until after 30–45 minutes before increasing from 35–45 minutes and peaking after more than 1.5 hours. These differences are directly attributable to the differences in PVS velocities and geometries: in the idealized models, the peak and average periarterial fluid velocities were approximately twice those of the vein, and the convective front thus appears twice as fast around the artery. The difference between the artery and the vein was greater in the image-based model, due to faster transport along the artery and slower transport along the vein.

### Tracer patterns depend on time of measurement

In Figures 7–8, we visualize the concentration field (above a given threshold) at the PVS outlets at a series of time points. After around 20 minutes, tracer had started to appear around the artery in the idealized model (Figure 7, *t* = 27 min). The corresponding apparent structure of the periarterial space is two disjoint compartments, one on each side of the artery. However, after 60 minutes, the tracer was distributed in the entire PVS cross-section. Analogously, 70 minutes after injection tracers appeared around the idealized vein and also seem to reveal two disjoint avenues of transport. After 100 minutes, both the arterial and venous PVSs were filled at the cross-sectional area at the measurement site in the idealized models. For the image-based models, the trends were similar, but included asymmetry due to the asymmetric geometries (Figure 8). After 15–20 minutes, tracer appeared on both sides of the image-based periarterial space, while it took up to 90 minutes for the venous PVS to reach similar levels of concentration at the measurement site.

**Figure 7.**
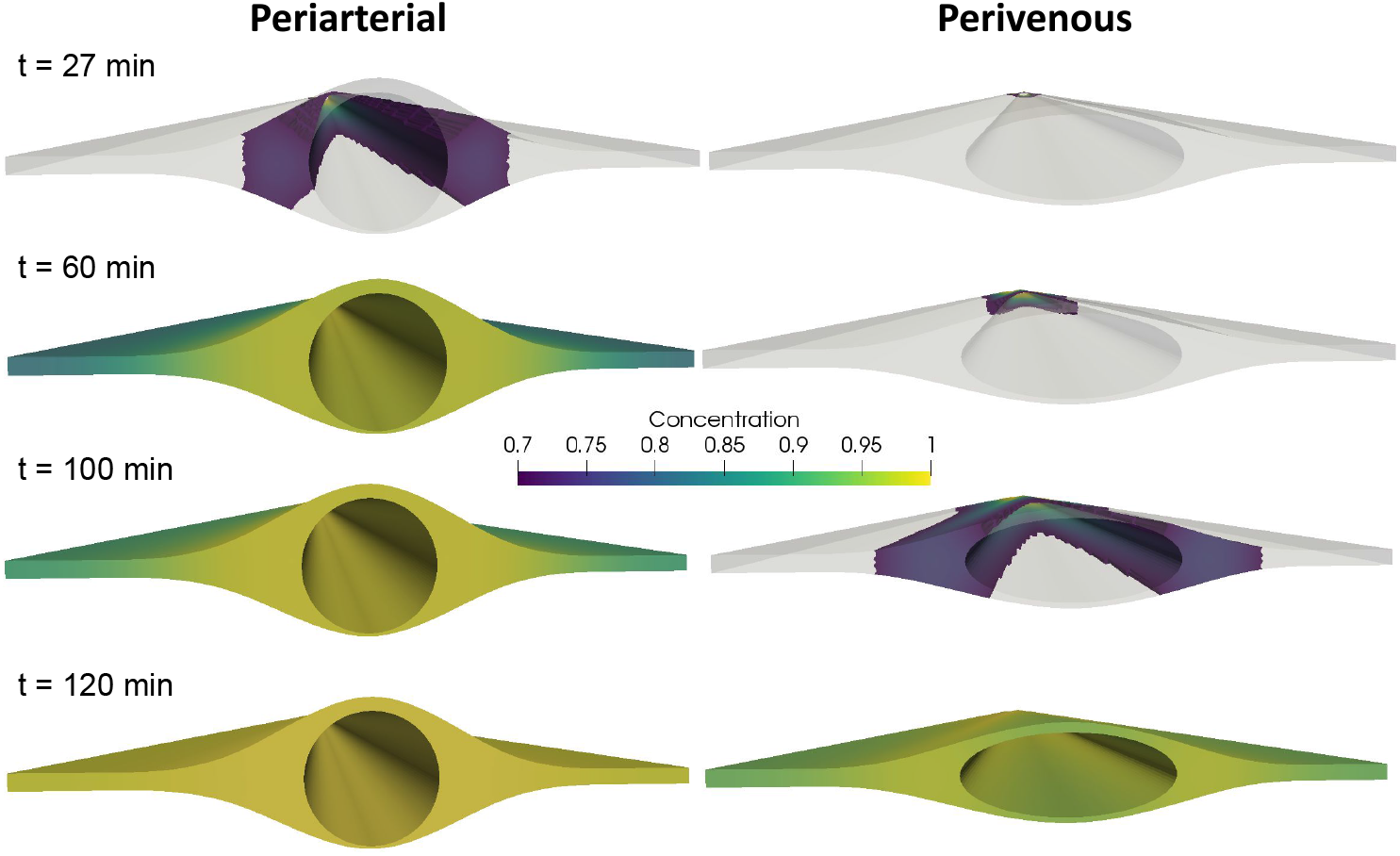
Tracer distribution around the artery (left) and the vein (right) at several time points after injection in the idealized geometries. The front end shows the site of measurement, while the point of injection is in the far end of the geometry. Colors indicate a threshold of 0.7. At 27 minutes (and at any time between 25 and 30 minutes) after injection, the areas of intense concentration around the artery suggest two disjoint PVS. After 60 minutes, the concentration is almost uniform at the cross section at the artery PVS, while no tracer is found around the vein. At 70 minutes after injection, tracer appears around the vein, apparently forming two disjoint PVS. After 100 minutes, tracer is distributed almost uniformly over both cross sections.

**Figure 8.**
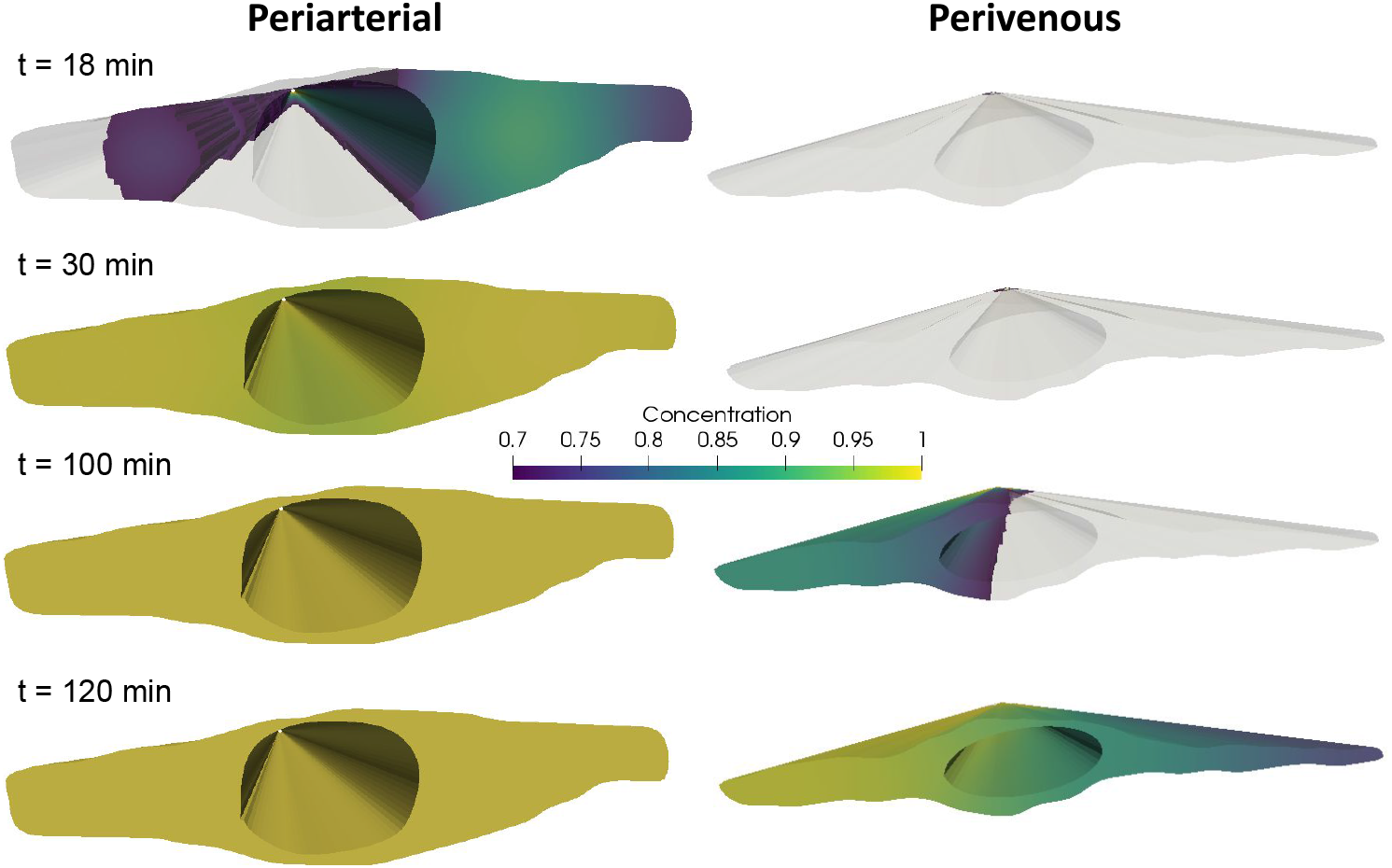
Tracer distribution around the artery (left) and the vein (right) at several time points after injection. The front end shows the site of measurement, while the point of injection is in the far end of the geometry. Colors indicate a threshold of 0.7. At 18 minutes (any time between 15 and 20 minutes) after injection, the areas of intense concentration around the artery suggest two disjoint PVS. After 90 minutes, the concentration is almost uniform at the cross section at the artery PVS, while the venous PVS is half covered, despite the fact that there is a small path for communication between the PVS on each side of the vessel. At 120 minutes after injection, tracer distributes almost uniformly around both the artery and the vein. In the extracted geometries we note asymmetry due to slightly larger PVS on one side of the vessel.

## Discussion

In this study, we used computational fluid dynamics to investigate differences in tracer transport along surface vessels of the brain. The geometrical models involved both idealized and image-based vessels. In both cases, we found higher PVS velocities in periarterial spaces compared to perivenous spaces. These differences were caused by the geometrical differences between the spaces and resulted in much faster transport along arteries than along veins. The surface vessels and PVSs modeled in this study should be distinguished from penetrating vessels and capillaries which may have different surroundings [31, 32].

In agreement with previous studies [15, 33], we show that for large molecules, such as e.g. 2000 kDa dextran, pure diffusion is too slow to explain the rapid tracer transport observed along the brain surface [7, 10]. Diffusive transport of the large tracers studied here occurs on the scale of days and months and not minutes or hours. Dispersion induced by arterial pulsations has been proposed as a mechanism to explain solute transport even in the absence of bulk flow [15, 33]. However, an increase by a factor of approximately two as demonstrated by Asgari et al. [15] does not seem sufficient to account for the rapid transport of large tracers in pial PVSs [7, 10]. By analytical models, Troyetsky et al. [23] have reached the same conclusion, namely that bulk flow velocities of ≈20 *µ*m/s are much more effective in transporting tracers than dispersion by arterial pulsations. On the other hand, within the brain parenchyma, transport of smaller tracers (e.g. <1 kDa gadubutrol) occur but slightly faster than by diffusion alone [34, 35, 36], and within a similar order of magnitude.

In our study, periarterial fluid velocities were but slightly lower than experimental evidence suggests (peak velocities of 11-23 *µ*m/s compared to ≈10-40 *µ*m/s) [7, 10]. Pulsatile flow patterns, not included in our models here, could further enhance tracer transport to some extent. According to data presented by Iliff et al. [11, Figure 4C,D], reduced vessel pulsations reduce tracer intensity at the measurement site by a factor close to 2. Local arterial wall pulsations can induce a small net flow, of 0.1 *µ*m/s over a 5 mm long PVS or up to 7 *µ*m/s in very long (100 mm) PVSs [16]. On the other hand, tracer infusions increase the intracranial pressure (ICP) [11], and may also give rise to a small but longlasting ICP gradient [37]. Yet, the experimentally observed rapid flows of macroparticles and large tracers in the pial PVSs do not seem to hinge upon such ICP increases [38]. A small static pressure gradient seems to be necessary for net flow of the reported magnitude [19, 16], yet the required gradient is of similar magnitude as naturally pulsatile pressure gradients within the brain [25].

In our simulations, tracers appeared first around arteries (after 5–20 minutes), then around veins (1–2 hours). These observations can be ascribed to the fact that PVS velocities were higher in the vicinity of arteries than around veins. According to the original glymphatic hypothesis [3], CSF flows in along arteries, CSF/ISF flows through the interstitium, and ISF is cleared along paravenous routes. The later appearance of tracers around veins on the brain surface would then be explained by this recirculation and clearance pathway. However, the delay observed by Iliff et al. [3] (0-30 min vs 1-2 hours for arteries vs veins) compares well with the results from our simulations for which no such recirculation was assumed. Instead, our data suggest that differences in periarterial and perivenous geometries explain the later appearance of tracer around veins, and not necessarily a full recirculation through a (glymphatic) interstitial pathway. We also note that, in agreement with our hypothesis, Ma et al. [39] observed spread of tracer directly from branches of the middle cerebral artery to some of the larger veins. In their study, it was concluded that tracer spread over the dorsal hemisphere was confined both to larger caliber arteries and veins at the brain surface [39].

Weller and coauthors [40] and many subsequent studies [41, 13, 15, 8, 14, 19, 16, 17] have suggested or assumed that pial PVSs take the form of annular or elliptic tubes that are separate(d) from the SAS, while Nedergaard et al. [7, 18] also suggest that the PVS forms disjoint compartments on each side of the blood vessels. We show here that thresholded representations of tracer concentration, and thus likely volumetric fluorescence images, will highlight areas with high bulk flow and essentially hide areas with low bulk flow if the measurement site is some distance away from the site of injection. More precisely, tracer images can, at some time points, display PVSs as separate both from the SAS and each other when they in fact form one continuous space (Figures 7–8) The effect of this gap on bulk flow may not be large, especially if the gap connecting two PVS around a vessel is small, but allows for diffusion and mixing of particles between the two spaces.

Imaging of the brain surface of humans and mice with OCT suggests that the PVS are widened sections of the SAS [10]. The round shape of arteries, which results from the relatively high intravascular pressure, may physically open up the space between the arachnoid and pia mater. Further away from vessels, the SAS may nearly collapse and not even be detectable. This could create the impression that PVS are separated from the SAS. From volumetric imaging of tracer transported via bulk flow, we show that the distinction between the two views might not be obvious. In both cases, more tracer will be seen in the large PVS areas on each side of the vessel, while there will be less, or even no tracer at all in other areas. In particular, our results demonstrate that tracer accumulation around vessels is not in disagreement with the theory that the PVS and SAS form one continuous space. In data provided by Schain et al. [8], the spread of tracer along PVS occurs rapidly, and travels ≈1 mm in a few minutes. Transport along the cortex surface perpendicular to the PVS however, is much slower and only covers a few *µ*m during the same time period, a transport more in line with diffusion given the size of tracer (3 kDa) used in their experiments [8]).

In terms of limitations of our study, we have not included arterial pulsations, brain compliance, or fluid or solute transport across the pial membrane. Arterial pulsations have been proposed to be the main driver of pulsatile PVS flow [11, 38] or even net flow [13, 7]. However, as recently pointed out by Troyetsky et al. [23], the bulk flow, and not the oscillatory movement, is the main mechanism of transport. As our simulated net flow was close to the experimentally observed range, we would expect to see but a minor additional effect of vessel wall pulsations or pulsatile pressure gradients on tracer transport. Similarly, by including brain compliance one could imagine a minor additional change in fluid velocities and dispersion [42].

Further, the OCT images used to generate the image-based geometries have relatively poor resolution compared to the size of the PVS. Yet, our findings show that any consistent difference in shape between arterial and venous PVS will lead to a change in transport velocity, and consequently a different time of appearance of tracer around arteries versus veins. We also note that we (by image quality) selected only two vessels to generate the image-based domains although we expect there to be large differences within both types of vessels (arteries and veins). The two extracted vessels were of different size, which may impact the size of the adjacent PVS. The images were extracted in 2D and extruded in the z-direction, and the PVS velocity was computed independent of the z-coordinate (velocity distribution is equal in all cross-sections of PVS). We thus omit bifurcations and the curvature of the vessel. However, as PVS flow has a very low Reynolds number (Re < 0.01), and as we have previously shown that bifurcations do not introduce complex flow patterns in the PVS [16], we therefore consider this simplification to be reasonably valid. The concentration threshold used to render volumetric images was arbitrarily set; a change in this value would not alter our conclusions, but would shift the arrival time of tracer shown in Figures 7–8.

In conclusion, we have shown that geometry differences between arterial and venous pial PVSs lead to higher net CSF flow, more rapid tracer transport and earlier arrival times of injected tracers in periarterial spaces compared to perivenous spaces. These findings are in agreement with experimentally observed differences in tracer arrival times, and can explain the rapid appearance of tracers around arteries, and the delayed appearance around veins without the need of a circulation through the parenchyma, but rather by direct transport along the PVS.

## Author’s contributions

V.V., M.E.R. conceived the experiments. V.V., M.E.R. and E.N.T.P.B wrote the paper. E.N.T.P.B provided OCT imaging data. All authors reviewed the manuscript.

## Acknowledgements

This study has received funding from the European Research Council (ERC) under the European Union’s Horizon 2020 research and innovation programme under grant agreement 714892. We would like to thank Dr. M. Almasian and Dr. B. Bedussi for their contribution to the OCT images.

## Availability of data and materials

The mesh-files used in this study can be found at the repository [21]. Additional files can be provided upon request to the corresponding author.

---

**Figure 9**\* **Additional file 1 (video)**: Perivascular flows resulting from a gradual change from periarterial to perivenous space.

**Figure 10**\* **Additional file 2 (video)**: Perivascular flows resulting from a gradual change in PVS widths.

**Figure 11**\* **Additional file 3 (figure)**: Numerical convergence study. Left: Concentration (AU) in idealized perivenous spaces vs time for different time resolutions ∆*t* = 10, 5, 2.5 s. Right: Concentration (AU) in idealized perivenous spaces vs time for different mesh resolutions (coarse, medium, fine). The refined meshes were constructed by uniform refinement of the original mesh. Simulation results are reported from the ‘medium’ mesh and the coarsest time step.

**Figure 12**\* **Additional file 4 (figure)**: The “medium” mesh resolution on geometries A0 and V0 used to perform the simulations

**Figure 13**\* **Additional file 5 (video)**: Concentrations in idealized periarterial and perivenous geometries. A threshold of 0.7 was used to visualize the transport of tracer. Tracers appear earlier around periarterial spaces due to higher convective velocities.

**Figure 14**\* **Additional file 6 (video)**: Concentrations in image-based periarterial and perivenous geometries. A threshold of 0.7 was used to visualize the transport of tracer. Tracers appear earlier around periarterial spaces due to higher convective velocities.

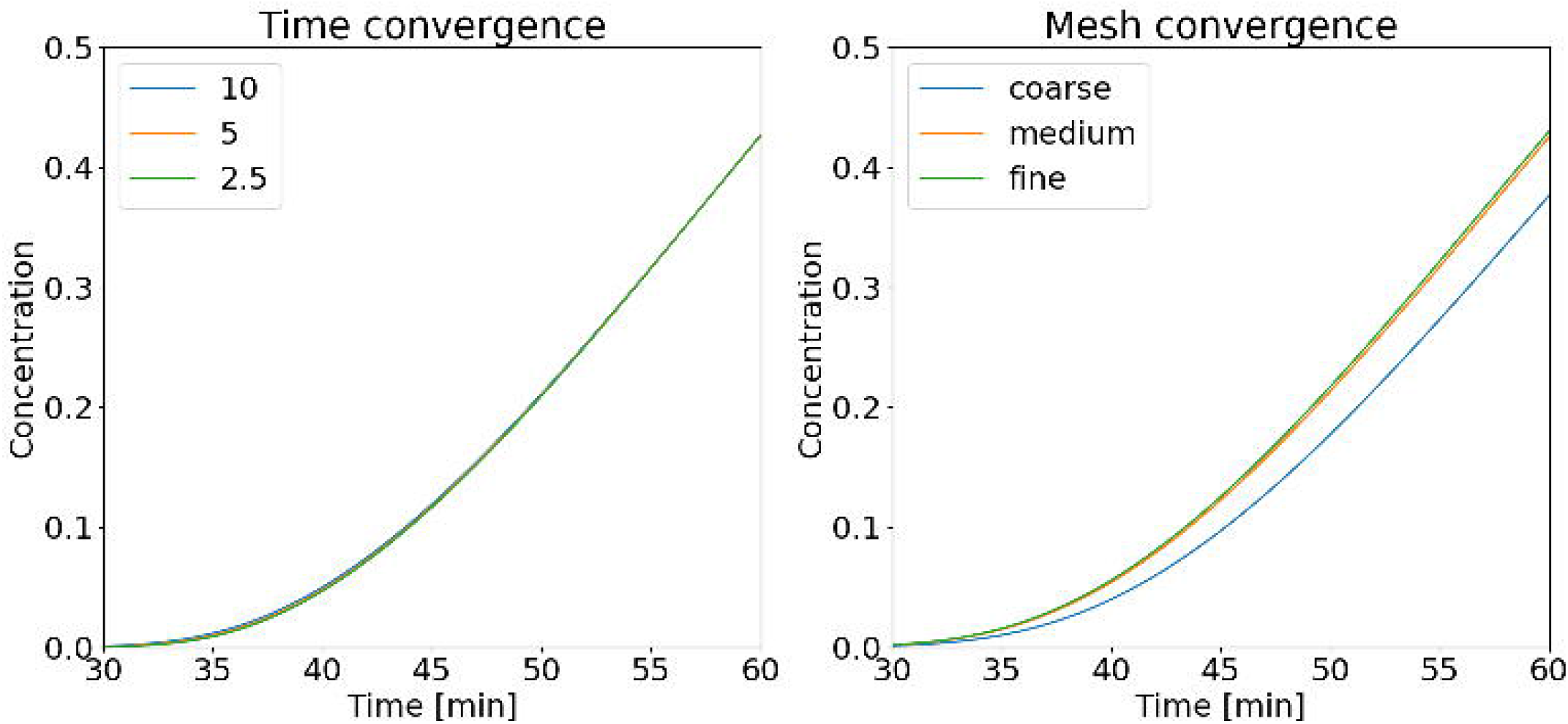

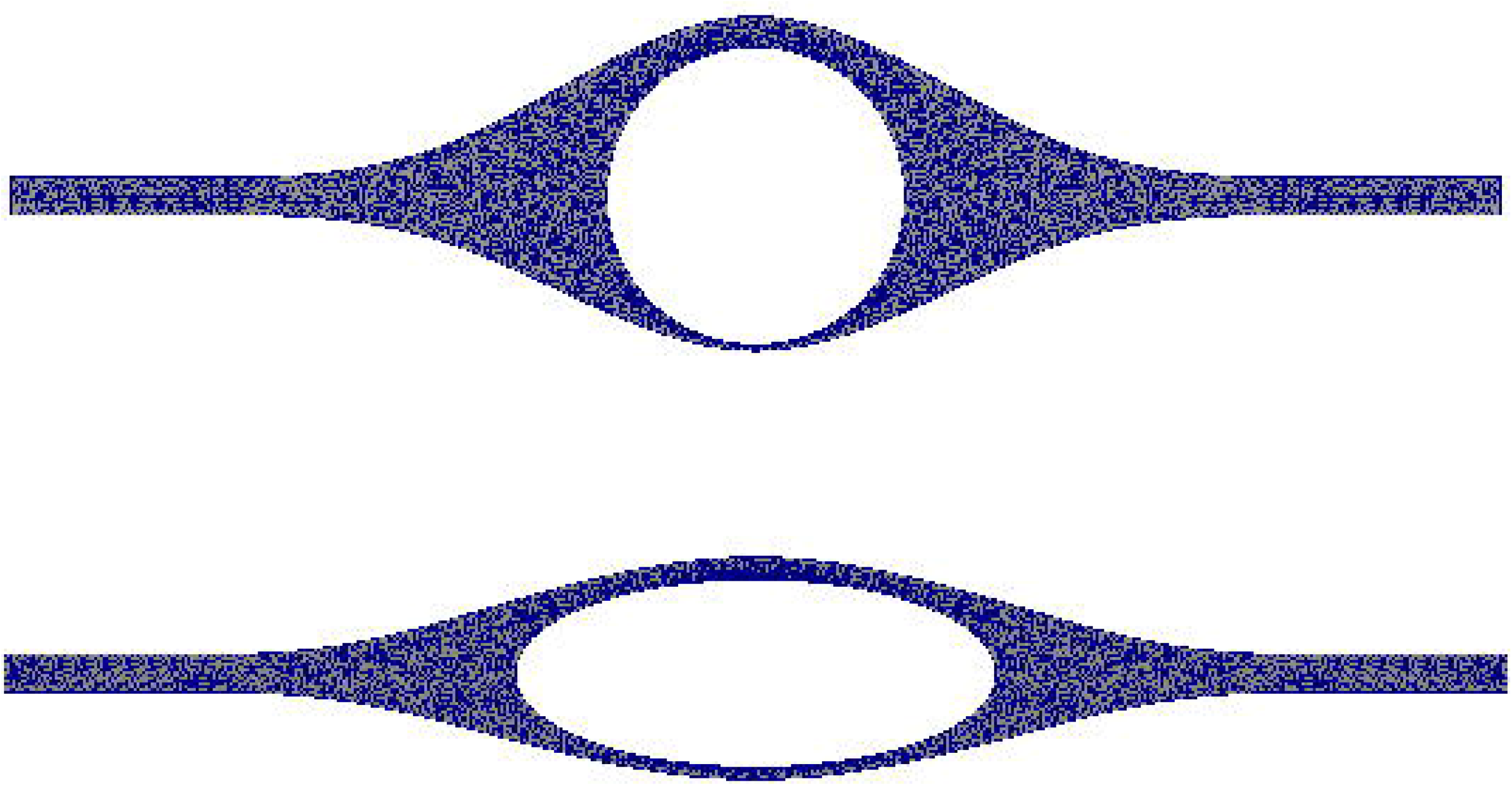

